# Chirped Speech (Cheech) Enables Rapid Assessment of Multi-Level Auditory Evoked Potentials During Speech-in-Noise Recognition

**DOI:** 10.64898/2026.08.19.745831

**Authors:** May Chao, C. Alise Holloway, Lee M. Miller, Kelsey Mankel

## Abstract

Difficulties understanding speech in noise remain a common complaint even among listeners with normal hearing sensitivity, highlighting the need for objective, more effective measures of real-world listening. The goal of this study was to validate the use of a novel, chirped-speech (Cheech) stimulus—continuous, naturally-spoken speech fused with chirps designed to elicit robust auditory evoked potentials—to characterize relationships between speech recognition, listening effort, and auditory neural encoding. Twenty-five normal-hearing adults completed a sentence-recognition task using both original (unmodified) and Cheech-modified AzBio sentence lists in quiet, +3 dB, and −3 dB signal-to-noise ratio (SNR) conditions while neural responses from the brainstem through cortex were recorded simultaneously. Speech recognition remained near ceiling in quiet but declined with decreasing SNR for both original and Cheech stimuli. Compared with clean speech, Cheech-modified speech showed slightly poorer recognition performance as SNR decreased and somewhat higher perceived effort overall. Yet, Cheech was highly effective at evoking auditory responses from the brainstem (auditory brainstem response, ABR) through the cortex (including middle- and late-latency responses, MLR and LLR) even with <5 minutes listening time per condition. Neural responses showed reduced amplitudes and prolonged latencies as SNR decreased. In general, ABR latencies and wave I amplitudes were associated with speech-in-noise recognition performance, whereas cortical responses (MLR Na, Nb, and LLR P1) were associated with subjective workload. These findings show that Cheech-modified speech preserves intelligibility while yielding robust, multilevel neural recordings during sentence perception, offering a promising approach to examine hierarchical auditory processing under ecologically relevant speech-in-noise conditions.

## 1 Introduction

Successful speech-in-noise perception relies on the brain’s ability to accurately encode complex acoustic signals amidst competing sounds. However, it remains unclear how different levels of the auditory pathway collectively contribute to an individual’s capacity to understand continuous speech in either quiet or noisy conditions. This gap highlights the need for approaches that can bridge traditional measures of auditory neural activity with real-world speech perception. The aim of the present study is to validate a novel method for measuring neural responses to continuous speech stimuli and to characterize neural biomarkers associated with individual speech perception abilities.

Common stimuli to assess speech perception abilities include meaningful monosyllabic words (e.g., NU-6, W-22), spondees, nonsense syllables (e.g., CUNY-NST), and sentences (e.g., AzBio, HINT, BKB-SIN, QuickSIN) (Lucks Mendel, 2025). Sentence-based tests in noise are considered to better represent real-world listening situations. For example, the Quick Speech-in-Noise (QuickSIN) test is a widely used clinical tool and provides an estimate of signal-to-noise ratio (SNR) loss (Billings et al., 2023; Killion et al., 2004). The AzBio Sentence Test is another example; its sentences are spoken by multiple talkers in a conversational style with limited contextual cues, making it effective for assessing speech understanding in more natural and challenging listening environments (Schafer et al., 2012; Spahr et al., 2012). Although these behavioral speech tests capture how well an individual recognizes speech, they provide limited insight into the underlying factors that influence performance. For instance, they cannot explain why two listeners with similar audiograms may perform differently in noisy or complex environments.

Listening under adverse conditions increases cognitive effort, which can in turn influence speech perception performance. Measures of listening effort and fatigue can therefore provide supplementary information about the cognitive resources required for speech (-in-noise) perception (Peelle, 2018). Subjective tools such as the NASA Task Load Index (NASA-TLX) allow individuals to report their perceived mental workload during listening tasks (Hart & Staveland, 1988; Mankel et al., in press). The NASA-TLX has been widely used to assess listening effort during speech perception tasks, with greater perceived workload often associated with increased difficulty understanding speech in noise (Bologna et al., 2013; Dimitrijevic et al., 2019; Nguyen et al., 2024; Seeman & Sims, 2015). However, these measures do not provide objective evidence to explain why performance varies across individuals, nor do they identify which stages of the auditory pathway underline these differences. Therefore, in addition to subjective measures of mental workload and listening fatigue, objective measures are needed to assess auditory pathway function and provide a more comprehensive understanding of speech perception.

Auditory evoked potentials (AEPs, also known as auditory event-related potentials, ERPs) can assess neural processing from lower (peripheral and brainstem) to higher (cortex) levels of the auditory pathway through short-latency (auditory brainstem response; ABR), middle-latency (MLR), and late-latency (LLR) responses, respectively. Among these measures, the ABR is considered the clinical “gold standard” for objective assessment of auditory neural function and is the most widely used electrophysiologic test in audiology practice (Hall, 2015a, 2015b; Mankel et al., 2026; The Joint Committee on Infant Hearing, 2019). However, this approach has two main limitations: (1) higher-level neural processes—particularly midbrain (MLR) and cortical (LLR) contributions—are rarely evaluated in routine clinical practice, and (2) the most common and effective transient stimuli used to evoke the ABR (e.g., clicks, tone bursts, and chirps) do not adequately reflect the neural encoding of naturally spoken speech (Brodbeck & Simon, 2020; Ding & Simon, 2014; Hamilton & Huth, 2020).

Recent research has increasingly focused on using continuous speech stimuli to elicit neural responses, enabling a more naturalistic assessment of the auditory system. These studies generally follow two distinct strategies: analysis-driven approaches, using techniques like temporal response functions (TRFs) to map stimulus features to estimated neural activity (Bachmann et al., 2021; Ding & Simon, 2014; Kulasingham et al., 2024; Maddox & Lee, 2018; Saiz-Alía & Reichenbach, 2020) and stimulus-modification approaches, which incorporate enhanced acoustic features such as glottal pulses fused with chirps or altered phase characteristics to increase the robustness of auditory evoked potentials (Backer et al., 2019; Mankel et al., in press; Polonenko & Maddox, 2021; Shan & Maddox, 2025; Shehabi et al., 2025; Stoll et al., 2025). Within these frameworks, there is a growing emphasis on incorporating measurements of both the subcortical and cortical responses to capture higher-level auditory processing beyond the brainstem (e.g., Bachmann et al., 2021; Mankel et al., in press; Shan et al., 2024; Shehabi et al., 2025).

Building on these advances, the “chirped-speech” (Cheech) technique integrates narrowband chirps with naturalistic speech to rapidly evoke responses from the brainstem to the cortex, allowing simultaneous measurement of multiple neural responses under ecologically valid listening conditions (Backer et al., 2019; Mankel et al., in press; Miller et al., 2020; Shehabi et al., 2025). Using short stories in mono- and dual-talker conditions, Cheech has recently revealed relationships between AEPs and measures of speech-in-noise processing, including selective attention, comprehension, and listening effort. Specifically, faster and/or more robust Cheech ABRs were associated with better target word identification accuracy and reaction times, narrative comprehension, and lower effort scores, highlighting Cheech’s potential utility for uncovering relationships between the brain and individual listening abilities (Mankel et al., in press). This foundation is supported by broader evidence indicating that neural encoding across the hierarchy (from subcortical phase-locking to cortical timing) is critical for speech perception in challenging environments (Bramhall et al., 2015; DiNino et al., 2025; Papakonstantinou et al., 2011). Despite these findings, most continuous-speech paradigms have relied on nonstandard narrative stimuli rather than standardized speech materials. Examining how multi-level neural responses are recorded during validated sentence-recognition tasks is a critical step toward translating laboratory findings into clinically meaningful tools for understanding individual differences in speech-in-noise listening abilities.

In this study, we aim to validate Cheech as a technique for assessing speech recognition by comparing original (unmodified) and Cheech-modified AzBio sentences. While simultaneously measuring Cheech-evoked neural responses at brainstem and cortical levels, we aim to characterize the effects of noise on neural encoding and establish brain–behavior relationships across the auditory hierarchy. We anticipate that the Cheech modification may pose a slight increase to listening task demands, but it will remain highly intelligible and provide a sensitive, multi-level view of individual speech perception abilities. Because more efficient neural processing (e.g., shorter latencies and higher synchrony) better facilitates the segregation of speech from noise (Bajo & King, 2012; Peelle, 2018), we hypothesize that faster, more robust neural encoding across the auditory pathway will be associated with enhanced speech-in-noise performance and reduced perceived effort, especially within the brainstem (Mankel et al., in press). Additionally, in light of evidence suggesting cortical responses evoked by continuous speech are sensitive to various cognitive factors (Shehabi et al., 2025), we predict cortical activity will be more strongly associated with listening effort and mental task load. Our study will be the first to apply the Cheech paradigm with standardized clinical speech materials to bridge the gap between electrophysiologic assessments and real-world speech understanding.

## 2 Methods

### 2.1 Participants

Twenty-five participants were recruited for this study (19 females, 6 males). To be eligible, individuals were required to be between 18 and 40 years old, right-handed, speak English as their primary language, and report no prior neurological or psychiatric conditions that could interfere with the goals of the study (i.e., speech perception, attention, or electrophysiological recordings such as epilepsy, stroke, ADHD, scalp lesions, etc.). The average age of participants was 23.0 years (SD = 2.65, range = 19–28). Participants reported an average of 17.1 years of education (SD = 2.22, range = 13–22) and 5.2 years of formal music training (SD = 6.70, range = 0–25). Socioeconomic status (SES) was estimated based on parents’ highest level of education using a 6-point scale, with an average score of 3.8 (SD = 1.08, range = 2–6, corresponding to high school diploma or GED through doctoral degree [e.g., PhD, MD, JD, EdD, ThD], respectively). All participants self-identified as right-handed and completed the Edinburgh Handedness Inventory to confirm handedness (mean laterality quotient = 0.99 indicating strongly right handed, SD = 0.02, range = 0.92–1.00; Oldfield, 1971). All participants demonstrated normal hearing, defined as air conduction thresholds ≤25 dB HL across octave frequencies from 250 to 8000 Hz in both ears. Participants were screened using the Quick Speech-in-Noise (QuickSIN) test to ensure they exhibited no more than a mild degree of signal-to-noise ratio (SNR) loss averaged across two lists (i.e., <7 dB SNR loss; mean = −0.46, SD = 2.27; Killion et al., 2004). The experiments conducted adhered to a protocol approved by the University of Memphis Institutional Review Board. Prior to participation, all subjects provided informed consent and were compensated for their time.

### 2.2 Stimuli

#### 2.2.1 AzBio stimuli

Auditory stimuli for this experiment were sourced from AzBio sentence lists, which were originally developed to assess speech perception abilities in individuals with hearing impairments and cochlear implant users. AzBio lists are comprised of 20 sentences spoken by 4 talkers (5 sentences each). Each list contains high- and low-context sentences crafted to isolate hearing ability by minimizing contextual linguistic influence (Spahr et al., 2012). To ensure consistent task difficulty across conditions, we followed the approach of Schafer et al. (2012) who validated the equivalency of the 15 AzBio sentence lists in noise for listeners with normal hearing (NH) as well as cochlear implant (CI) users. For NH participants, lists 1, 6, 7, 12, and 14 resulted in significantly lower or higher average performance compared to the 10 remaining lists (Schafer et al., 2012). We thus excluded these lists from our study. The remaining 10 lists (2, 3, 4, 5, 8, 9, 10, 11, 13, 15) were used across three listening conditions: quiet, +3 dB SNR, and –3 dB SNR (three lists per condition), including one list reserved for practice. Pilot testing indicated that three lists per condition was sufficient to produce clear, robust neural responses by Cheech (see Cheech stimuli below). Multi-talker babble from the original AzBio audio test was adapted for the two speech-in-noise conditions.

#### 2.2.2 Cheech stimuli

To facilitate the simultaneous recording of the auditory brainstem (ABR), middle latency (MLR), and late latency responses (LLR) while participants completed a speech recognition task, the AzBio sentence lists were modified using chirped-speech (Cheech), a process designed to enhance neural synchrony to continuous speech stimuli (Backer et al., 2019; Miller et al., 2020; Shehabi et al., 2025; Mankel et al., in press). Cheech blends natural, continuous speech with upward frequency-modulated chirp stimuli designed to evoke more synchronized brainstem responses than traditional clicks by compensating for traveling wave delays along the basilar membrane (Backer et al., 2019; Dau et al., 2000; Elberling et al., 2007).

In the Cheech synthesis, segments of glottal pulse energy generated by vocal fold vibrations during voiced speech were partially replaced with synthetic chirp energy according to previously defined protocols (Backer et al., 2019; Mankel et al., in press; Miller et al., 2020; Shehabi et al., 2025) using custom MATLAB code (MathWorks, Inc.; vR2022a). Chirps were time-locked to glottal pulses and inserted only during segments of sufficient voiced power.

These segments were identified from filtered audio (20-1000 Hz) when the speech envelope between 20-40 Hz surpassed approximately 28% of the overall root-mean-square (RMS) amplitude for periods at least 50 ms long (i.e., normalized amplitude voicing threshold of 0.03). Timing of glottal pulses during voiced periods were determined through a speech resynthesis process based on the TANDEM-STRAIGHT toolbox (Kawahara et al., 1999, 2008; Kawahara & Morise, 2011).

After glottal pulses were identified, chirp trains were created with precise temporal alignment to the timing of the glottal pulses with a minimum spacing of 18.2 ms between chirps (55 Hz). The inter-stimulus interval of the chirps varied with the natural fluctuations in continuous speech, but certain constraints were added to optimize simultaneous measurement of multiple auditory evoked responses. The first chirp in a voiced segment was always followed by a minimum of 48 ms before the second chirp in a sequence, then additional chirps were added through the maximum voicing duration of 400 ms, after which 50 ms gaps were inserted between chirps to prevent very long chirp trains (e.g., during loud speech modulations). Chirps were normalized to approximately 0.1× the overall speech RMS amplitude per each AzBio sentence list. All AzBio audio files were normalized prior to the Cheech synthesis procedure to ensure equivalent chirp amplitudes relative to speech levels across sentence lists.

The original audio was re-filtered into alternating, octave-wide frequency bands from 0-250, 500-1000, 2000-4000, and 11,000-∞ Hz (10^th^-order Butterworth filter). The chirp trains were constrained to energy within the interleaved frequency bands, specifically 250-500, 1000-2000, and 4000-11,000 Hz (4^th^-order Butterworth filter). Finally, the alternating chirp and speech bands were added together to produce the Cheech-ed AzBio audio. A comparison of the original and Cheech-modified speech is shown in Figure 1.

**Figure 1:**
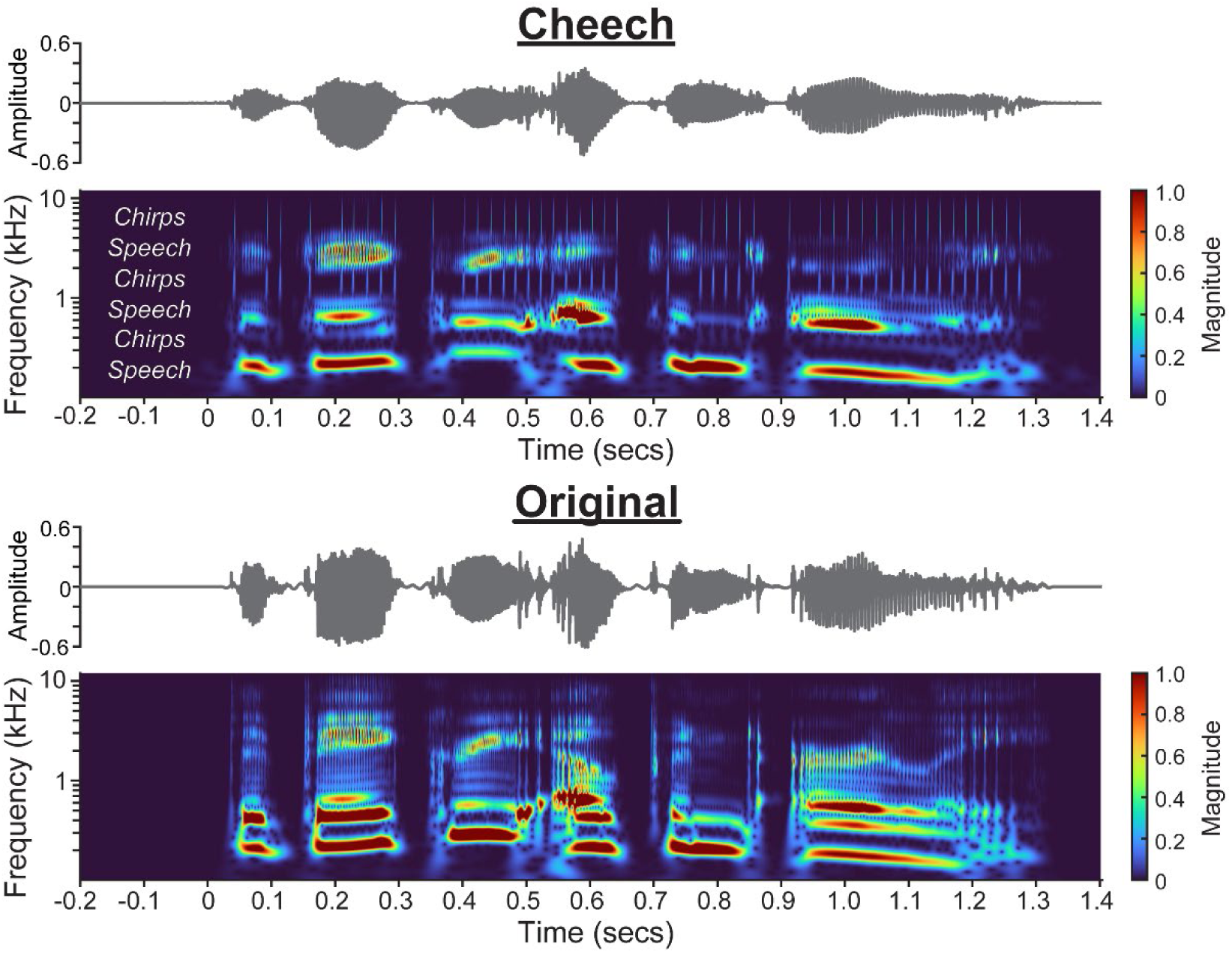
Cheech-modified (top) and original (clean) speech waveforms (bottom). Cheech modifies AzBio sentences by replacing glottal pulse energy in predefined frequency bands with temporally aligned chirps, interleaving speech and chirp frequency bands while preserving global speech structure and intelligibility. The stimulus is designed to enhance neural synchrony for simultaneous recording of ABR, MLR, and LLR. See 2.2.2 Cheech stimuli for details. The stimulus for both is AzBio list 9, sentence 12: “The bakery opened early”.

### 2.3 Subjective measures

To assess perceived task-related effort and workload during the speech recognition task, participants completed a modified, computerized version of the NASA Task Load Index (NASA-TLX; Hart & Staveland, 1988). Task load was assessed by six questions evaluating mental demand, physical demand, temporal demand, performance, effort, and frustration. The computerized version featured prompts with sliding scales labeled at the extremes (e.g., very low to very high), and participants used the keyboard to enter their subjective ratings. Responses were converted into numerical values from 0 to 100, where higher scores indicated greater task load or effort. The total NASA-TLX score reflects the average across the six questions. Additionally, a seventh question was included to evaluate the evolution of task difficulty over the course of each block, but this data is not included in the present analysis.

### 2.4 Procedure

The audio signal was routed through a Tucker-Davis Technologies (TDT) RP2 processor at a sample rate of 48,828 Hz and calibrated to an average sound level of approximately 60 dBA (slow weighting), measured with a Larson-Davis sound level meter. Multi-talker babble noise was added as needed to create varying SNR conditions. Stimulus presentation and EEG triggering was managed using a custom MATLAB script that interfaced with TDT hardware via RPvdsEx software.

Participants were seated comfortably in an electrically shielded, sound-attenuating booth. Auditory stimuli were presented binaurally via electromagnetically shielded insert earphones (3M E-A-RTONE 3A) equipped with disposable foam tips. Three lists were played for each of the three conditions (quiet, +3 dB, and −3 dB SNR) and repeated twice: once as the original, unmodified speech, and once as Cheech-ed audio (3 SNR conditions × 3 lists × 2 = 18 total lists/blocks). These lists were presented in a pseudo-randomized order, ensuring that the same SNR condition did not appear back-to-back and the same list did not appear within 5 lists of each other to minimize learning effects. Presentation order was also counterbalanced across “forward” and “backwards” sequences to reduce potential order-fatigue effects—half the participants heard the 18 lists in one order, and the other half heard the lists played in the reversed order (i.e., the last list played in the “forward” order was the first list played in the “backward” order).

Participants were asked to listen to auditory stimuli and verbally repeat the speech they heard. The original AzBio audio file timing was preserved for each list, where gaps between each sentence provided sufficient time for the participant to respond. Verbal responses were recorded at a sampling rate of 44.1 kHz using Audacity and a Realistic 33-1070B omnidirectional microphone routed through a Focusrite Scarlett 18i20 audio interface. The microphone was placed within one foot from the subject and adjusted as needed to ensure consistent audio quality. Input levels were maintained around −12 dB to avoid clipping. Speech recognition accuracy was scored by two trained raters, either in real-time or offline. Final scores were averaged across the two raters. Interrater reliability between the two scorers was very good (ICC(2,1) = 0.89, 95% CI = 0.87-0.91), indicating strong agreement across all scored items. After completing each sentence list, the monitor displayed questions from the computerized version of the NASA-TLX to assess perceived task-related effort (Hart & Staveland, 1988). Each block lasted ∼3-5 minutes. The total experimental session duration was approximately two hours, with rest breaks provided as needed to minimize fatigue.

### 2.5 EEG recordings

Continuous EEG data were recorded using a Neuroscan SynAmps 2/RT amplifier and Curry 7 software (Compumedics, Inc.). Signals were digitized at a sampling rate of 10 kHz with a single-channel configuration. The active (non-inverting) electrode was positioned at Fz with linked mastoid reference electrodes at M1 (left mastoid) and M2 (right mastoid) and the ground electrode placed at Fpz. Electrode impedances were maintained below 3 kΩ across all channels to ensure signal quality. Participants were instructed to remain relaxed during recording.

#### 2.5.1 EEG preprocessing and analysis

EEG data were preprocessed in MATLAB 2022a (MathWorks Inc.) using a combination of EEGLAB functions (Delorme & Makeig, 2004), ERPLAB functions (Lopez-Calderon & Luck, 2014), and custom MATLAB scripts. Continuous data were first filtered with an eighth-order Butterworth filter using response-specific frequency bands: 100-1500 Hz for auditory brainstem responses (ABR), 15-200 Hz for middle latency responses (MLR), and 0.5-40 Hz for late latency responses (LLR). Epoch windows were defined by response type: −2-15 ms for ABRs, −5-60 ms for MLRs, and −50-500 ms for LLRs^1^. AEP waveforms were averaged across trials for each participant and condition. The averaged responses were baseline corrected according to each pre-stimulus window. Artifact rejection was performed; epochs with deflections exceeding ± 150 uV for ABR and MLRs and ± 250 uV for LLRs were excluded from the AEP averages (mean % total rejected trials ± SD: ABR 0.04 ± 0.001%, MLR 0.10 ± 0.002%, LLR 3.20 ± 0.069%.

The ERPLAB toolbox was used to measure amplitude and latency values of local maximum/minimum peaks within each time window (over 3 consecutive sample points) (Lopez-Calderon & Luck, 2014). If no local peak was found, the absolute voltage within the time window was recorded. AEP components were then measured using predefined latency windows specific to each response type and polarity. Measurement time windows were based on previously established Cheech protocols (Backer et al., 2019; Mankel et al., in press; Shehabi et al., 2025) and tweaked as needed to optimally measure the correct peaks according to visual inspection of the grand average and individual subject waveforms. For the ABR, previous evidence suggests peak-to-trough measures provide a more robust estimate of Cheech-evoked brainstem activity (Mankel et al., in press). ABR components therefore included wave I (positive peak, 2-4.5 ms), wave III (positive peak, 4.5-6.75 ms), and wave V (positive peak, 6.75-10 ms) as well as the subsequent negative troughs for wave I (3.4-5 ms), wave III (5.7-7.5), and wave V (8.4-11.5).

Given their role in neurodiagnostic measurements (Hall III, 2007), inter-peak latencies (IPLs) between ABR components were also calculated for I-III, III-V, and I-V IPLs. MLR components were analyzed as P0 (positive peak, 10-16 ms), Na (negative peak, 12-25 ms), Pa (positive peak, 20-35 ms), Nb (negative peak, 30-45 ms), and Pb (40-59.5 ms). LLR component time windows include P1 (positive peak, 40-90 ms) and N1 (mean amplitude, 110-200 ms). The mean amplitude measurement provides a better estimate of N1 since Cheech LLRs are typically characterized by a sustained negativity after P1 (Backer et al., 2019; Shehabi et al., 2025). All latency measurements were then adjusted to correct for a 1 ms transducer delay as well as a 0.3 ms delay between the Cheech-chirps and accompanying EEG triggers (total = 1.3 ms)^2^.

### 2.6 Statistical analysis

Statistical analyses were performed in SAS (SAS Institute, Inc., v3.82). A significance level of α = 0.05 was used for all analyses. Mixed-effects linear regression models were employed to analyze the data using PROC GLIMMIX. The models included a random subject intercept to account for inherent variability across individuals and improve generalizability of the results. Behavioral outcomes such as speech recognition accuracy and subjective ratings of task effort were modeled as dependent variables. For behavioral-only tests, the main fixed effects tested included stimulus condition (Cheech versus clean speech), signal-to-noise ratio (quiet, +3 dB, and −3 dB SNR), and their interaction. Since the Cheech stimuli were specifically designed to elicit robust auditory evoked potentials, only Cheech responses were included for analyses with the neural data. EEG metrics were used both as outcome variables (for brain-only tests, with SNR condition as the main fixed effect) and as predictors in brain-behavior analyses (including SNR condition and their interaction). As noted in the results, all participants scored above 90% speech recognition for the quiet condition. Thus, Cheech-in-quiet was excluded from the brain-behavior analyses (speech recognition only) to facilitate model convergence and prevent violation of regression model assumptions regarding normality and homogeneity of residual variance. If any interaction terms were not significant, they were removed and the simplified model results were reported. Unless otherwise specified, Tukey corrections were applied to post hoc pairwise comparisons. Observations with conditional studentized residuals exceeding ± 3 were marked as outliers. If removing an outlier observation (i.e., entire subject for that measurement) did not alter statistical conclusions, the original results prior to outlier removal were reported; else, any statistics with outliers excluded are noted in the results.

## 3 Results

### 3.1 Behavioral data

#### 3.1.1 Speech recognition scores vary by conditions (clean vs. Cheech) and SNR

We analyzed the effects of the Cheech modification, signal-to-noise ratio condition (SNR), and their interaction on behavioral speech recognition (Figure 2A). Mixed effects regression results included significant main effects of SNR (F_2,120_ = 1725.65, p < .0001), clean versus Cheech (F_1,120_ = 397.09, p < .0001), as well as an interaction of the two variables (F_2,120_ = 90.85, p < .0001) on AzBio speech recognition. Tukey-adjusted post-hoc tests indicated comparable ceiling-level performance for both clean speech and Cheech (t_120_ = −1.25, p = .2129), suggesting that Cheech speech remained intelligible, with a slight drop in Cheech recognition performance for the +3 dB (t_120_ = −13.17, p < .0001) and −3 dB SNR conditions (t_120_ = −20.10, p < .0001) compared to the original AzBio sentences. Additionally, while Cheech recognition performance decreased with decreasing SNR (all post hoc p’s < .05), AzBio recognition for the unmodified sentence lists was similar between quiet and +3 dB SNR (albeit indicating a similar decreasing trend; t_120_ = −2.23, p = .0700).

**Figure 2:**
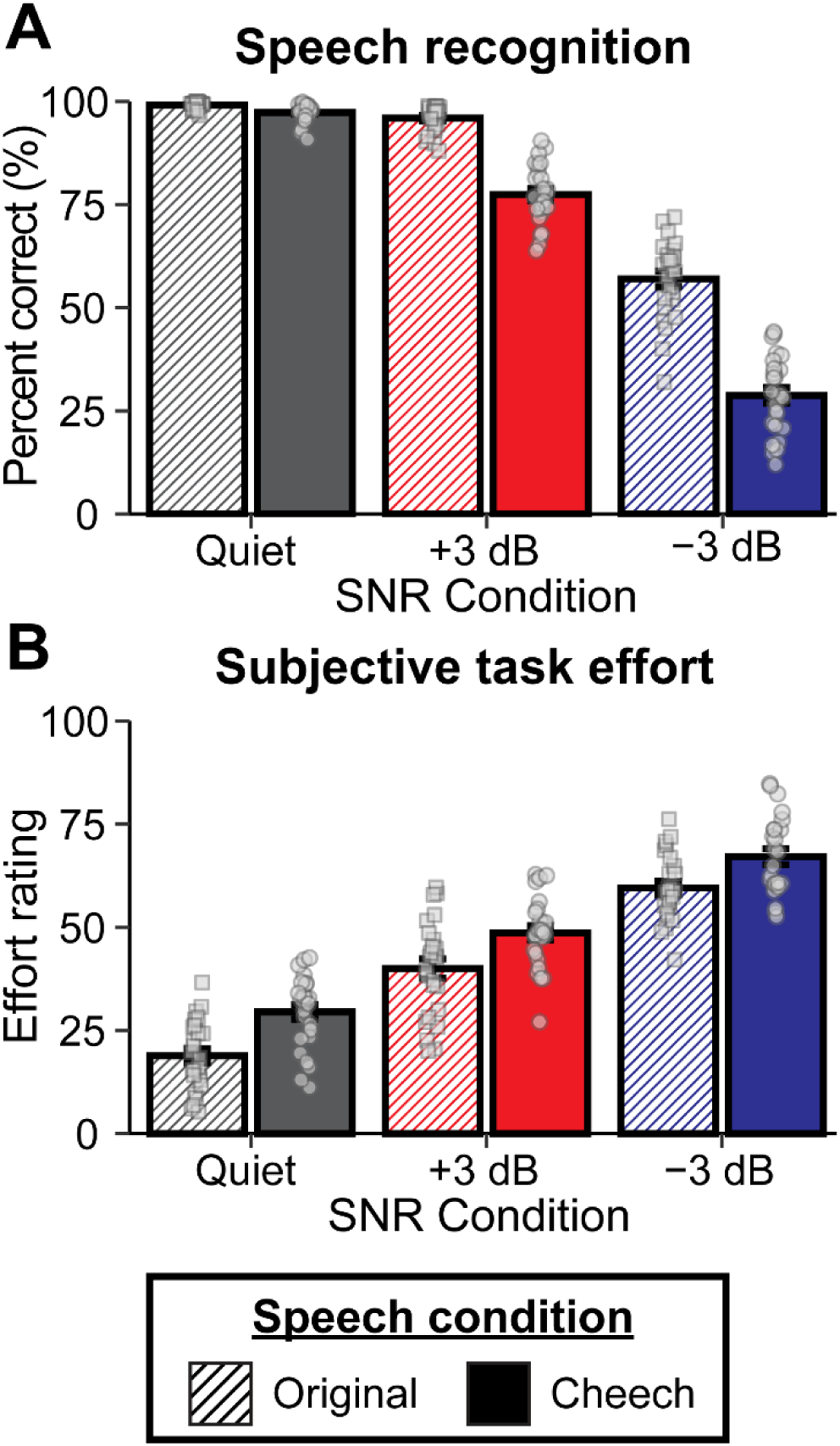
(A) Speech recognition differs between Cheech and clean speech, and this effect varies across SNR conditions (worse Cheech-listening performance at lower SNRs). (B) Subjective task effort (NASA Task Load Index, NASA-TLX) differs between Cheech and clean speech, with higher reported effort for Cheech overall and increased effort at lower SNRs. Bars = mean +/- standard error.

#### 3.1.2 Higher subjective task load ratings (i.e., greater workload/listening effort) reported for Cheech and lower SNRs

Similar analyses were conducted to evaluate the effects of Cheech, SNR, and their interaction on subjective task effort according to average NASA-TLX ratings (Hart & Staveland, 1988; Figure 2B). Subjects reported greater task load for Cheech compared to clean speech (F_1,122_ = 59.15, p < .0001) and with decreasing SNR (overall F_2,122_ = 376.64, p < .0001; all post hoc pairwise p’s < .0001). The interaction term was not significant.

We also examined whether subjective ratings were associated with recognition performance. A 3-way interaction was observed for subjective effort, SNR, and clean versus Cheech (F_2,114_ = 4.46, p = .0136), so we conducted follow-up analyses on Cheech and clean audio separately. Only main effects of effort ratings and SNR were observed for Cheech-modified AzBio. That is, higher effort ratings were associated worse Cheech recognition performance (F_1,47_ = 5.79, p = .0201). SNR effects on recognition performance were consistent with those reported above. However, the two-way interaction between task effort and SNR was significant for clean speech (F_2,45_ = 12.56, p < .0001) in addition to a main effect of task effort (F_1,45_ = 14.15, p = .0005). In general, higher effort scores were associated with worse speech recognition, but this effect was stronger for −3 dB SNR compared to the quiet (t_45_ = 4.22, p = .0001) and +3 dB conditions (t_45_ = 4.74, p < .0001).

### 3.2 Electrophysiological data

Since Cheech is designed to elicit robust auditory event-related potentials (ERPs) in the context of continuous speech, we evaluated the effects of SNR on the auditory brainstem, middle latency, and late latency responses for the Cheech-modified AzBio sentences (ABR, MLR, and LLR, respectively). As expected, decreasing SNR was generally characterized by decreased amplitudes and increased latencies across the auditory hierarchy (Figure 3). This effect was significant for ABR wave V amplitudes and latencies (overall condition effect, latencies: F_2,48_ = 20.28, p < .0001; with one outlier excluded, amplitudes: F_2,46_ = 19.31, p < .0001). The quiet condition was associated with larger wave V amplitudes and faster latencies than +3 dB (latencies: t_48_ = −3.3090, p = .0050; amplitudes: t_46_ = 2.7897, p = .0205) and −3 dB SNR (latencies: t_48_ = −6.3673, p < .0001; amplitudes: t_46_ = 6.2046, p < .0001). +3 dB, in turn, was associated with larger wave V amplitudes and faster latencies than −3 dB SNR (latencies: t_48_ = −3.0583, p = .0100; amplitudes: t_46_ = 3.4149, p = .0038). Compared to the −3 dB SNR condition, Cheech-in-quiet was also associated with shorter III-V IPLs (t_48_ = −3.2203, p = .0064) and larger wave I amplitudes (with one outlier excluded, t_46_ = 2.4709, p = .0446). Additionally, MLR Na peak amplitudes were largest for Cheech-in-quiet compared to +3 and −3 dB SNR (with one outlier excluded, t_46_ = −3.8986, p = .0009 & t_46_ = −4.7172, p < .0001, respectively). No other pairwise comparisons significantly differed across SNR conditions and ERP components.

**Figure 3:**
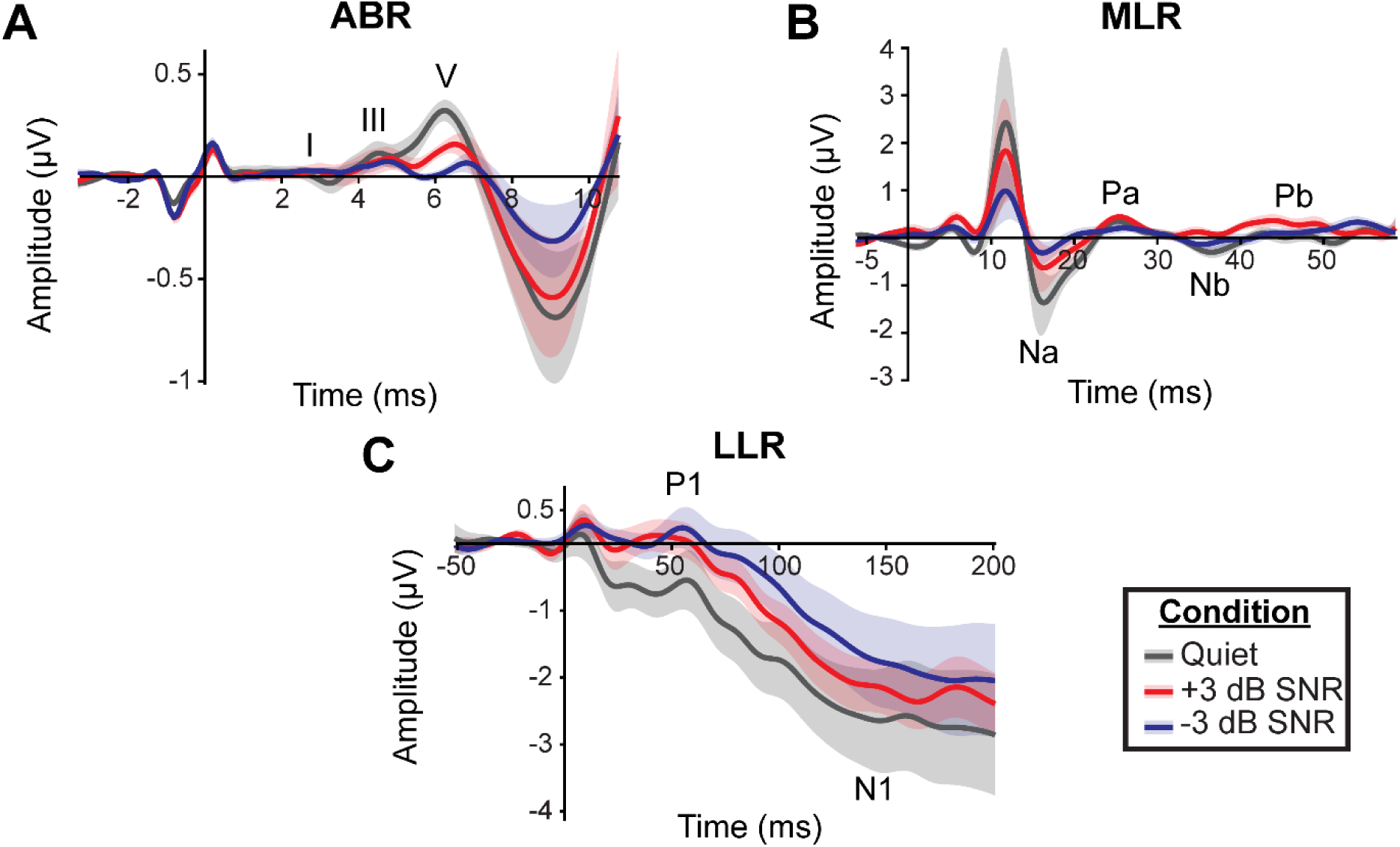
Auditory evoked potentials (AEPs) to Cheech-modified AzBio sentences across SNR conditions. (A) ABRs, (B) MLRs, and (C) LLRs generally exhibit smaller amplitudes and delayed latencies at lower SNRs.

**Figure 4:**
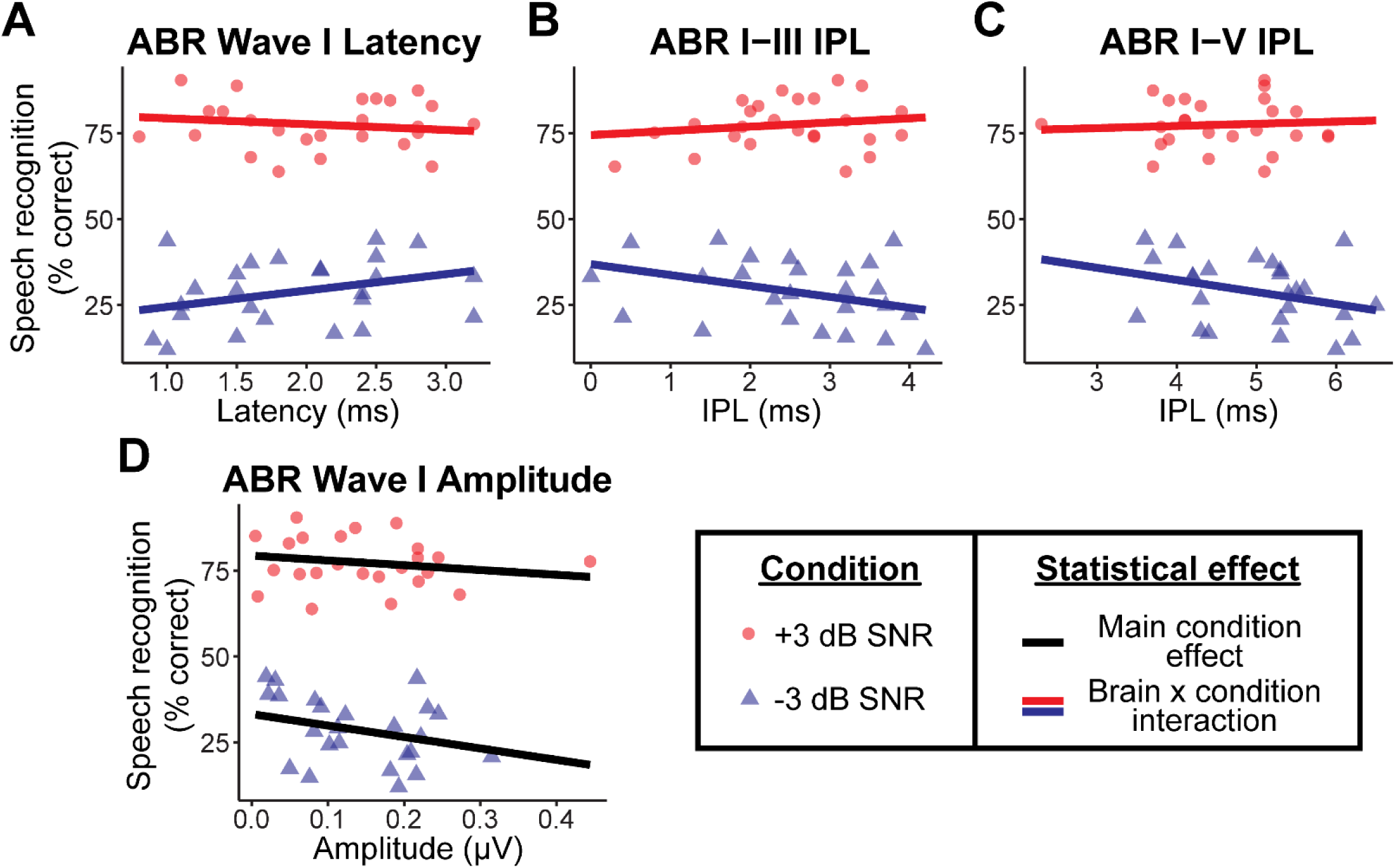
Relationships between ABR measures and Cheech speech-in-noise recognition. (A-C) Longer wave I latencies and shorter I-nd I-V inter peak latencies (IPL) were associated with poorer recognition performance, particularly in the −3 dB SNR condition pared to +3 dB SNR. (D) Smaller wave I amplitudes were also associated with better overall speech-in-noise recognition.

**Figure 5:**
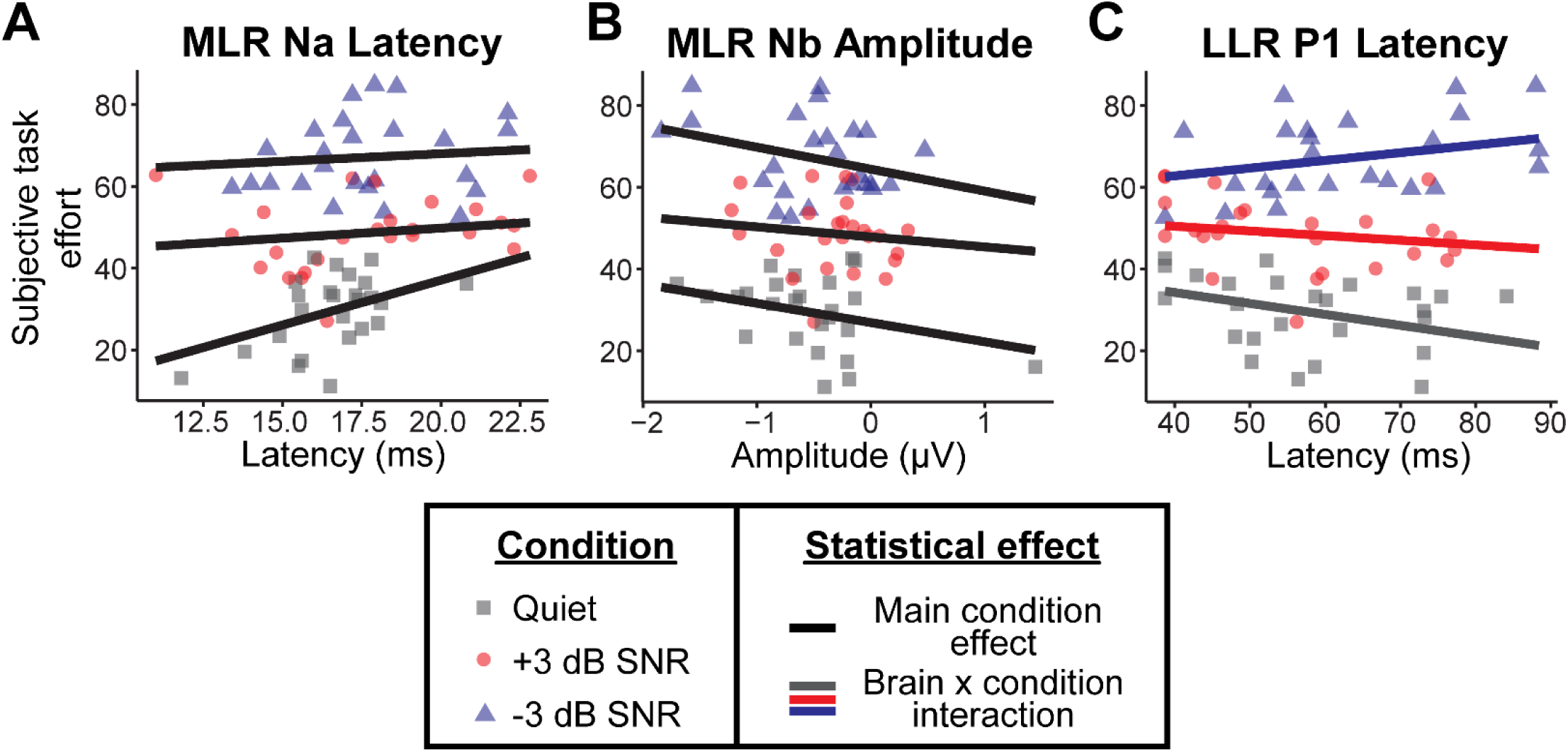
Relationships between neural measures and subjective task effort. (A) Delayed Na latencies were associated with higher rted effort. (B) Larger (i.e., more negative) Nb amplitudes were also linked to greater effort. (C) Faster LLR P1 peak latencies in t compared to −3 dB SNR were associated with lower effort ratings.

### 3.3 Relationships between brain responses and speech recognition

One advantage of our Cheech technique is the ability to measure auditory brain responses from the brainstem and cortex simultaneously while participants complete a behavioral listening task (e.g., Mankel et al., in press; Shehabi et al., 2025). We thus wanted to characterize the relationship between neural encoding along the auditory pathway and speech (Cheech) recognition performance in our study. As shown in Figure 2 and noted in 2.6 Statistical analysis, performance was at ceiling for the Cheech-in-quiet condition. The limited variability across subjects makes characterizing the brain-behavior relationship for this condition difficult.

Consequently, our analyses here focus on Cheech-in-noise behavioral and neural responses (i.e., +3 and −3 dB SNR). Of our ERP measures, the ABR was most strongly associated with speech-in-noise recognition performance. An interaction was observed between SNR condition and wave I peak latencies (F_1,22_ = 6.69, p =.0168), wave III amplitudes (F_1,22_ = 4.99, p =.0359), I-V IPL (F_1,22_ = 4.82, p = .0389), and I-III IPL (F_1,22_ = 7.24, p = .0134). We then evaluated marginal effects to reveal how ABR measures related to behavior for each SNR condition individually. In the harder −3 dB SNR condition, results indicated a relationship between ABR latencies and speech-in-noise recognition (Bonferroni adjusted, wave I peak latencies: t_22_ = 2.78, p = .0218; I-V IPL: t_22_ = −2.53, p = .0381; I-III IPL: t_22_ = −3.22, p = .0078) and a trend for wave III peak amplitudes (Bonferroni adjusted, t_22_ = −2.35, p = .0560). Specifically, better speech recognition performance in the harder −3 dB SNR condition was characterized by slower wave I latencies and, in turn, shorter I-III and I-V IPLs. Similar effects were not observed in the +3 dB SNR condition (wave I latencies: t_22_ = −.96, p = .6946; wave III amplitudes: t_22_ = .69, p = .9974; I-V IPL: t_22_ = .44, p = 1; I-III IPL: t_22_ = .99, p = .6657). Additionally, smaller wave I amplitudes were associated with better Cheech-in-noise recognition overall (main effect F_1,22_ = 4.60, p = .0432 with non-significant interaction term included; F_1,23_ = 2.31, p = 0.1423 with interaction term removed). Collectively, our results suggest that lower-level neural encoding of auditory signals, particularly at the brainstem, contributes to individual speech in noise recognition abilities.

### 3.4 Relationships between brain responses and subjective effort ratings

We also investigated potential neural factors associated with subjective listening effort in our Cheech paradigm. An interaction between SNR condition and LLR P1 peak latencies was observed for NASA-TLX ratings (F_2,45_ = 4.40, p = .0180). Faster latencies for quiet compared to −3 dB SNR was associated with lower effort ratings (t_45_ = −2.87, Bonferroni-adjusted p = .0372).

Additionally, main effects of Na peak latencies and Nb amplitudes indicated that delayed Na responses (main effect F_1,45_ = 5.09, p = .0290 with non-significant interaction term included; F_1,47_ = 2.91, p = .0948 with interaction term removed) and larger, more negative Nb amplitudes were associated with higher effort ratings overall (F_1,47_ = 5.69, p = .0211). Thus, higher-level auditory cortical generators appear linked with subjective perceptions of listening task effort.

## 4. Discussion

The present study examined how Cheech-modified speech and varying signal-to-noise ratios (SNRs) influence speech recognition, subjective listening effort, and auditory neural encoding. The Cheech paradigm permitted simultaneous acquisition of auditory evoked potentials from the brainstem through the cortex while participants completed standard AzBio sentence recognition, providing a unified view of behavioral and neural processing under different listening demands. Overall, listeners showed near-ceiling performance in quiet but reduced recognition and increased perceived effort as SNR decreased, with Cheech speech eliciting slightly greater listening demands than clean speech. Across the auditory neuroaxis, decreasing SNR produced smaller amplitudes and delayed latencies. Auditory brainstem latency measures were most strongly associated with speech-in-noise performance whereas cortical responses were more closely related to subjective listening effort.

Speech recognition performance for both clean and Cheech-modified AzBio sentences remained near ceiling in the quiet condition (99.09 ± 1.35% and 97.32 ± 2.65% mean and standard deviation, respectively) but somewhat diverged as different SNR levels were introduced. The ceiling performance in quiet suggests that these changes do not significantly affect speech understanding, at least for our subjects who presented with normal hearing. This finding supports the work of Backer et al. (2019), demonstrating that the Cheech stimulus remains a promising paradigm when applied to speech testing. To our knowledge, this is the first study to apply the Cheech modification standardized clinical stimuli and conduct auditory evoked potential (AEP) and speech testing simultaneously. The divergence in scores across SNR conditions suggests that Cheech modification does impact intelligibility in noisy backgrounds compared to original, unmodified speech. According to previous studies with similar participant characteristics, the 77.45% Cheech recognition accuracy in +3 dB SNR (18.50% drop compared to clean speech) and 28.76% accuracy in −3 dB SNR (28.24% drop) corresponds to roughly ∼3-5 dB SNR loss to achieve equivalent performance on original, unmodified AzBio speech recognition (Holder et al., 2018; Schafer et al., 2012). This range approximately falls within a mild SNR loss range as suggested by the QuickSIN test, for example (Killion et al., 2004; Killion & Niquette, 2000). Accounting for this SNR difference or recalibrated norms would be needed if the Cheech versions were to be deployed as a clinical AzBio test in noise. However, our results collectively suggest that Cheech is highly effective at assessing speech recognition both in quiet and noisy backgrounds.

The NASA Task Load (TLX) scores suggested greater perceived effort both as the noise level increased and for Cheech compared to original speech. These results match our expectation that both background noise and the Cheech modifications increase perceptual demands (Pichora-Fuller et al., 2016). Even when participants understood much of the speech, they reported exerting more listening effort than they did for clean speech. Cheech may make perceptual tasks more difficult by reducing the clarity of the speech signal, especially in background noise. The specific design of our study may have also influenced effort ratings, where the original AzBio audio in quiet condition provided a clean baseline (as indicated by a mean NASA-TLX score <20 out of 100) whereby any subsequent alterations where judged as more effortful, whether background noise or altered by Cheech.

The neural data showed a similar SNR-dependent pattern across the auditory hierarchy. Decreasing SNR was associated with reduced amplitudes and/or delayed latencies in ABR, MLR, and LLR responses, indicating that noisier listening conditions degrade encoding at multiple stages of the auditory pathway. This finding is consistent with the view that continuous speech can reveal neural responses from brainstem to cortex under ecologically relevant conditions (Brodbeck & Simon, 2020; Maddox & Lee, 2018; Polonenko & Maddox, 2021). It also supports the rationale for using a chirp-based speech stimulus: by enhancing the ability to elicit robust responses, Cheech makes it possible to assess multiple auditory generators simultaneously during a single listening task.

The relationship between neural encoding and behavioral outcomes suggests that different stages of the auditory hierarchy contribute to distinct aspects of the listening experience. Our results indicate that ABR components are the primary predictors of speech-in-noise recognition, particularly in the challenging −3 dB SNR condition. This is consistent with established models showing that degraded subcortical encoding is linked to poorer speech-in-noise performance (Bramhall et al., 2015; Papakonstantinou et al., 2011; Saiz-Alía et al., 2019). Specifically, better recognition scores were associated with slower wave I peak latencies and shorter inter-peak latencies in our study. Given that wave I reflects the initial peripheral encoding at the level of the auditory nerve, these findings underscore the importance of early neural precision for successful top-down decoding. The finding that smaller wave I amplitudes were associated with better recognition warrants caution, however; amplitudes exhibit substantial inter-individual variability in normal-hearing listeners and is strongly influenced by anatomical factors (e.g., head size, skull thickness), physiology (e.g., outer hair cell function, cochlear synaptic integrity), and recording parameters (e.g., electrode montage, stimulus intensity), rather than directly reflecting perceptual ability (Bramhall, 2021; Verhulst et al., 2016). In contrast, latency and inter-peak latency reflect neural synchrony and phase-locking precision, aspects of subcortical timing that have been linked to different aspects of speech listening performance (Mankel et al., in press). Yet, mixed results have been observed between transient-evoked ABR relationships and speech-in-noise perception (Bramhall et al., 2015; DiNino et al., 2025; Papakonstantinou et al., 2011). Collectively, the present results suggest that early encoding within the auditory pathway serves as a reliable indicator of individual perceptual differences, at least in the context of continuous speech recognition.

In contrast, middle- and late-latency potentials appear more closely linked to the subjective experience of the task rather than recognition accuracy. We observed that lower effort ratings were associated with a larger shift in P1 latencies between quiet and noise.

Furthermore, larger Nb amplitudes and delayed Na latencies were both linked to higher overall effort ratings on the NASA-TLX. These results suggest a functional distinction in the auditory pathway: while the brainstem manages the neural precision needed for recognition, the auditory cortex reflects the cognitive workload required to maintain attention during speech listening.

By combining Cheech with standardized AzBio sentence testing, we demonstrate a feasible, efficient method to measure auditory neural responses while listeners perform a clinically meaningful speech task. This is a notable advantage because many recent continuous-speech EEG paradigms rely on non-standardized stimuli such as narratives and/or do not compare with behavioral performance measured at the same time (Bachmann et al., 2021; Ding & Simon, 2012; Etard et al., 2019; Kulasingham et al., 2024; Maddox & Lee, 2018; Mankel et al., in press; Polonenko & Maddox, 2021; Saiz-Alía et al., 2019; Shan & Maddox, 2025; Shehabi et al., 2025; Teoh et al., 2022). In contrast, AzBio is already familiar in clinical practice and is designed to assess speech understanding in both quiet and noise (Holder et al., 2018; Schafer et al., 2012; Spahr et al., 2012). Our single-channel electrode montage, consistent with acquisition setups commonly used for ABR screening and diagnostic testing, further supports translational application by demonstrating feasibility of a standard, simple, and cost-effective EEG recording setup for continuous-speech electrophysiology. This makes our paradigm especially promising for future work aimed at bridging laboratory measures and clinical assessment.

These results also have potential relevance for listeners who report difficulty understanding speech in noise despite normal audiometric thresholds. Standard audiograms and even traditional speech scores in quiet often fail to capture subtle deficits in speech processing that emerge only under more challenging listening conditions (Beck et al., 2018; Billings et al., 2023; Killion & Niquette, 2000; Musiek et al., 2017; Peelle, 2018; Pienkowski, 2017; Spankovich et al., 2018; Tremblay et al., 2015). A paradigm like Cheech-AzBio-in-noise may help identify which listeners show poorer neural encoding—and where along the auditory pathway—greater listening effort, or weaker speech-in-noise performance even when traditional clinical measures appear normal (DiNino et al., 2025; Shehabi et al., 2025). In that sense, the method may eventually contribute to a more individualized understanding of auditory function and help guide patient-centered assessment (Bramhall et al., 2015; Papakonstantinou et al., 2011). More broadly, this approach supports the idea that clinically relevant speech testing combined with novel electrophysiological techniques can reveal meaningful differences in how listeners process speech in everyday listening environments.

Several limitations should be considered. First, the sample consisted of listeners with normal hearing and relatively good speech-in-noise performance, which may have reduced behavioral variability and limited the range of detectable brain–behavior relationships. Second, the observed relationships should be interpreted cautiously because experimental paradigms, stimulus modifications, recording variability, and small sample sizes can all influence behavioral and neural results. Future work addressing these issues will be important for determining the potential clinical-translational applicability of Cheech stimuli and similar continuous speech-based approaches.

### 4.1 Conclusion

Cheech-modified speech was highly effective at simultaneous measurement of both behavior and neural responses in the context of a sentence-based recognition task, with listeners showing lower speech recognition and higher perceived effort at more challenging SNRs. Specifically, ABR components were strongly linked to speech-in-noise performance, suggesting that lower-level auditory processing at the brainstem influences speech perception abilities in noise. Cortical responses, meanwhile, were more closely tied to perceived workload and listening effort. Collectively, these findings provide valuable insight into how both lower- and higher-level auditory processes work together to support real-world speech understanding. Our study further highlights the advantage of simultaneously measuring neural responses while completing a standardized speech test. This paradigm may help identify and differentiate listeners who struggle with speech-in-noise, offering a potential solution for future clinical assessment.

## 5 Conflicts of interest

Lee M. Miller is an inventor on intellectual property related to chirped-speech (Cheech) owned by the Regents of University of California, not presently licensed.

## 6 Declaration of generative AI and AI-assisted technologies

Generative AI tools (ChatGPT) were used in a limited capacity for this work, primarily assisting with minor text editing and basic phrasing. All significant ideas, drafting, analysis, and editing were carried out by the authors. The authors take full responsibility for the content of the published article.

## Acknowledgements

The authors are grateful to our team members Jill Dodson, Amy Cox, and Mira Milman for their contributions to data collection and organization as well as Daniel C. Comstock for his advice during study design and analysis. This work was supported by the Center for Research Initiatives and Strategies for the Communicatively Impaired (CRISCI) at the University of Memphis. Data used for the current study are available from the corresponding author on reasonable request.

## Footnotes

1 Epoch windows were defined prior to the 1.3 ms latency correction (transducer delay of 1 ms + chirp-trigger delay of 0.3 ms). Establishing epoch windows prior to the latency shift allowed us to define baseline correction values from a pre-stimulus period before the onset of an electromagnetic artifact generated by the triggering equipment, most clearly evident in the ABR waveforms (artifact onset t=0ms prior to the latency shift or −1.3 ms after the latency shift; Figure 3A). Epoch windows after the latency shift were: −3.3-13.7 ms for ABRs, −6.3-58.7 for MLRs, and −51.3-498.7 ms for LLRs.

2 Measurement windows after accounting for the 1.3 ms latency shift would be: wave I 0.7-2.2 ms, post-wave I trough 2.1-3.7 ms, wave III 3.2-5.45 ms, post-wave III trough 4.3-6.2 ms, wave V 5.45-8.7 ms, post-wave V trough 7.1-10.2 ms, P0 8.7-14.7 ms, Na 10.7-23.7 ms, Pa 18.7-33.7 ms, Nb 28.7-43.7 ms, Pb 38.7-58.2 ms, P1 38.7-88.7 ms, and N1 108.7-198.7 ms.

## References

Bachmann, F. L., MacDonald, E. N., & Hjortkjær, J. (2021). Neural Measures of Pitch Processing in EEG Responses to Running Speech. Frontiers in Neuroscience, 15, 738408. 10.3389/fnins.2021.738408

Backer, K. C., Kessler, A. S., Lawyer, L. A., Corina, D. P., & Miller, L. M. (2019). A novel EEG paradigm to simultaneously and rapidly assess the functioning of auditory and visual pathways. Journal of Neurophysiology, 122(4), 1312–1329. 10.1152/jn.00868.2018

Bajo, V. M., & King, A. J. (2012). Cortical modulation of auditory processing in the midbrain. Frontiers in Neural Circuits, 6, 114. 10.3389/fncir.2012.00114

Beck, D. L., Danhauer, J. L., Abrams, H. B., Atcherson, S. R., Brown, K., Chasin, M., Clark, J. G., Placido, C. D., Edwards, B., Fabry, D. A., Flexer, C., Fligor, B., Frazer, G., Galster, J. A., Gifford, L., Johnson, C. E., Madell, J., Moore, D. R., Roeser, R. J., … Wolfe, J. (2018). Audiologic Considerations for People with Normal Hearing Sensitivity Yet Hearing Difficulty and/or Speech-in-Noise Problems. Hearing Review, 11.

Billings, C. J., Olsen, T. M., Charney, L., Madsen, B. M., & Holmes, C. E. (2023). Speech-in-Noise Testing: An Introduction for Audiologists. Seminars in Hearing, 45(1), 55–82. 10.1055/s-0043-1770155

Bologna, W. J., Chatterjee, M., & Dubno, J. R. (2013). Perceived listening effort for a tonal task with contralateral competing signals. The Journal of the Acoustical Society of America, 134(4), EL352–EL358. 10.1121/1.4820808

Bramhall, N. F. (2021). Use of the Auditory Brainstem Response for Assessment of Cochlear Synaptopathy in Humans. The Journal of the Acoustical Society of America, 150(6), 4440. 10.1121/10.0007484

Bramhall, N. F., Ong, B., Ko, J., & Parker, M. (2015). Speech Perception Ability in Noise is Correlated with Auditory Brainstem Response Wave I Amplitude. Journal of the American Academy of Audiology, 26(5), 509–517. 10.3766/jaaa.14100

Brodbeck, C., & Simon, J. Z. (2020). Continuous speech processing. Current Opinion in Physiology, 18, 25–31. 10.1016/j.cophys.2020.07.014

Dau, T., Wegner, O., Mellert, V., & Kollmeier, B. (2000). Auditory brainstem responses with optimized chirp signals compensating basilar-membrane dispersion. The Journal of the Acoustical Society of America, 107(3), 1530–1540. 10.1121/1.428438

Dimitrijevic, A., Smith, M. L., Kadis, D. S., & Moore, D. R. (2019). Neural indices of listening effort in noisy environments. Scientific Reports, 9, 11278. 10.1038/s41598-019-47643-1

Ding, N., & Simon, J. Z. (2012). Emergence of neural encoding of auditory objects while listening to competing speakers. Proceedings of the National Academy of Sciences, 109(29), 11854–11859. 10.1073/pnas.1205381109

Ding, N., & Simon, J. Z. (2014). Cortical entrainment to continuous speech: Functional roles and interpretations. Frontiers in Human Neuroscience, 8. 10.3389/fnhum.2014.00311

DiNino, M., Crowell, J., Kloiber, I., & Polonenko, M. J. (2025). The relationship between auditory brainstem responses, cognitive ability, and speech-in-noise perception among young adults with normal hearing thresholds. Hearing Research, 460, 109243. 10.1016/j.heares.2025.109243

Elberling, C., Don, M., Cebulla, M., & Stürzebecher, E. (2007). Auditory steady-state responses to chirp stimuli based on cochlear traveling wave delay. The Journal of the Acoustical Society of America, 122(5), 2772–2785. 10.1121/1.2783985

Etard, O., Kegler, M., Braiman, C., Forte, A. E., & Reichenbach, T. (2019). Decoding of selective attention to continuous speech from the human auditory brainstem response. NeuroImage, 200, 1–11. 10.1016/j.neuroimage.2019.06.029

Hall III, J. W. (2007). New Handbook for Auditory Evoked Responses. Pearson.

Hall, J. W. (2015a). eHandbook of Auditory Evoked Responses: Principles, Procedures & Protocols.

Hall, J. W. (2015b). Update on Auditory Evoked Responses: Value of Chirp Stimuli in ABR/ASSR Measurement. Audiology Online. https://www.audiologyonline.com/articles/update-on-auditory-evoked-responses-17434

Hamilton, L. S., & Huth, A. G. (2020). The revolution will not be controlled: Natural stimuli in speech neuroscience. *Language*, Cognition and Neuroscience, 35(5), 573–582. 10.1080/23273798.2018.1499946

Hart, S. G., & Staveland, L. E. (1988). Development of NASA-TLX (Task Load Index): Results of Empirical and Theoretical Research. In Advances in Psychology (Vol. 52, pp. 139–183). Elsevier. 10.1016/S0166-4115(08)62386-9

Holder, J. T., Levin, L. M., & Gifford, R. H. (2018). Speech recognition in noise for adults with normal hearing: Age-normative performance for AzBio, BKB-SIN, and QuickSIN. *Otology & Neurotology: Official Publication of the American Otological Society*, American Neurotology Society [and] European Academy of Otology and Neurotology, 39(10), e972–e978. 10.1097/MAO.0000000000002003

Kawahara, H., Masuda-Katsuse, I., & de Cheveigné, A. (1999). Restructuring speech representations using a pitch-adaptive time–frequency smoothing and an instantaneous-frequency-based F0 extraction: Possible role of a repetitive structure in sounds. Speech Communication, 27(3), 187–207. 10.1016/S0167-6393(98)00085-5

Kawahara, H., & Morise, M. (2011). Technical foundations of TANDEM-STRAIGHT, a speech analysis, modification and synthesis framework. Sadhana, 36(5), 713–727. 10.1007/s12046-011-0043-3

Kawahara, H., Morise, M., Takahashi, T., Nisimura, R., Irino, T., & Banno, H. (2008). Tandem-STRAIGHT: A temporally stable power spectral representation for periodic signals and applications to interference-free spectrum, F0, and aperiodicity estimation. 2008 IEEE International Conference on Acoustics, Speech and Signal Processing, 3933–3936. 10.1109/ICASSP.2008.4518514

Killion, M. C., & Niquette, P. A. (2000). What can the pure-tone audiogram tell us about a patient’s SNR loss? The Hearing Journal, 53(3), 46–53.

Killion, M. C., Niquette, P. A., Gudmundsen, G. I., Revit, L. J., & Banerjee, S. (2004). Development of a quick speech-in-noise test for measuring signal-to-noise ratio loss in normal-hearing and hearing-impaired listeners. The Journal of the Acoustical Society of America, 116(4 Pt 1), 2395–2405. 10.1121/1.1784440

Kulasingham, J. P., Bachmann, F. L., Eskelund, K., Enqvist, M., Innes-Brown, H., & Alickovic, E. (2024). Predictors for estimating subcortical EEG responses to continuous speech. PLOS ONE, 19(2), e0297826. 10.1371/journal.pone.0297826

Lopez-Calderon, J., & Luck, S. J. (2014). ERPLAB: An open-source toolbox for the analysis of event-related potentials. Frontiers in Human Neuroscience, 8. 10.3389/fnhum.2014.00213

Lucks Mendel, L. (2025). Speech Audiometry. In Audiology Diagnosis (3rd ed., pp. 149–170). Thieme.

Maddox, R. K., & Lee, A. K. C. (2018). Auditory Brainstem Responses to Continuous Natural Speech in Human Listeners. eNeuro, 5(1). 10.1523/ENEURO.0441-17.2018

Mankel, K., Comstock, D. C., Bormann, B. M., Das, S., Sagiv, D., Brodie, H., & Miller, L. M. (in press). Auditory brainstem encoding of speech-in-noise predicts word identification, narrative comprehension, and subjective listening effort in a selective attention task. 2024.12.23.629710. 10.1101/2024.12.23.629710

Mankel, K., Price, C. N., Milman, M., & Austin, A. (2026). Survey of audiologists’ clinical practice patterns and perceptions of electrophysiologic assessments. American Journal of Audiology. 10.1044/2026_AJA-25-00334

Miller, L. M., Moore, B. D., & Bishop, C. W. (2020). Frequency-multiplexed speech-sound stimuli for hierarchical neural characterization of speech processing (United States Patent No. PCT/US15/40629). https://patents.google.com/patent/WO2016011189A1/en

Musiek, F. E., Shinn, J., Chermak, G. D., & Bamiou, D.-E. (2017). Perspectives on the Pure-Tone Audiogram. Journal of the American Academy of Audiology, 28(7), 655–671. 10.3766/jaaa.16061

Nguyen, D. L., Valentin, O., Lehmann, A., & Prévost, F. (2024). A Multimodal Investigation of Listening Effort in Single-Sided Deafness. American Journal of Audiology, 33(4), 1341–1349. 10.1044/2024_AJA-24-00073

Oldfield, R. C. (1971). The assessment and analysis of handedness: The Edinburgh inventory. Neuropsychologia, 9(1), 97–113. 10.1016/0028-3932(71)90067-4

Papakonstantinou, A., Strelcyk, O., & Dau, T. (2011). Relations between perceptual measures of temporal processing, auditory-evoked brainstem responses and speech intelligibility in noise. Hearing Research, 280(1), 30–37. 10.1016/j.heares.2011.02.005

Peelle, J. E. (2018). Listening Effort: How the Cognitive Consequences of Acoustic Challenge Are Reflected in Brain and Behavior. Ear and Hearing, 39(2), 204–214. 10.1097/AUD.0000000000000494

Pichora-Fuller, M. K., Kramer, S. E., Eckert, M. A., Edwards, B., Hornsby, B. W. Y., Humes, L. E., Lemke, U., Lunner, T., Matthen, M., Mackersie, C. L., Naylor, G., Phillips, N. A., Richter, M., Rudner, M., Sommers, M. S., Tremblay, K. L., & Wingfield, A. (2016). Hearing Impairment and Cognitive Energy: The Framework for Understanding Effortful Listening (FUEL). Ear and Hearing, 37, 5S. 10.1097/AUD.0000000000000312

Pienkowski, M. (2017). On the Etiology of Listening Difficulties in Noise Despite Clinically Normal Audiograms. Ear and Hearing, 38(2), 135–148. 10.1097/AUD.0000000000000388

Polonenko, M. J., & Maddox, R. K. (2021). Exposing distinct subcortical components of the auditory brainstem response evoked by continuous naturalistic speech. eLife, 10, e62329. 10.7554/eLife.62329

Saiz-Alía, M., Forte, A. E., & Reichenbach, T. (2019). Individual differences in the attentional modulation of the human auditory brainstem response to speech inform on speech-in-noise deficits. Scientific Reports, 9(1), Article 1. 10.1038/s41598-019-50773-1

Saiz-Alía, M., & Reichenbach, T. (2020). Computational modeling of the auditory brainstem response to continuous speech. Journal of Neural Engineering, 17(3), 036035. 10.1088/1741-2552/ab970d

Schafer, E. C., Pogue, J., & Milrany, T. (2012). List Equivalency of the AzBio Sentence Test in Noise for Listeners with Normal-Hearing Sensitivity or Cochlear Implants. Journal of the American Academy of Audiology, 23(7), 501–509. 10.3766/jaaa.23.7.2

Seeman, S., & Sims, R. (2015). Comparison of Psychophysiological and Dual-Task Measures of Listening Effort. Journal of Speech, Language, and Hearing Research, 58(6), 1781–1792. 10.1044/2015_JSLHR-H-14-0180

Shan, T., Cappelloni, M. S., & Maddox, R. K. (2024). Subcortical responses to music and speech are alike while cortical responses diverge. Scientific Reports, 14(1), 789. 10.1038/s41598-023-50438-0

Shan, T., & Maddox, R. K. (2025). Comparing methods for deriving the auditory brainstem response to continuous speech in human listeners. Imaging Neuroscience, 3, IMAG.a.19. 10.1162/IMAG.a.19

Shehabi, S., Comstock, D. C., Mankel, K., Bormann, B. M., Das, S., Brodie, H., Sagiv, D., & Miller, L. M. (2025). Individual Differences in Cognition and Perception Predict Neural Processing of Speech in Noise for Audiometrically Normal Listeners. eNeuro, 12(4). 10.1523/ENEURO.0381-24.2025

Spahr, A. J., Dorman, M. F., Litvak, L. M., Van Wie, S., Gifford, R. H., Loizou, P. C., Loiselle, L. M., Oakes, T., & Cook, S. (2012). Development and validation of the AzBio sentence lists. Ear and Hearing, 33(1), 112–117. 10.1097/AUD.0b013e31822c2549

Spankovich, C., Gonzalez, V. B., Su, D., & Bishop, C. E. (2018). Self reported hearing difficulty, tinnitus, and normal audiometric thresholds, the National Health and Nutrition Examination Survey 1999-2002. Hearing Research, 358, 30–36. 10.1016/j.heares.2017.12.001

Stoll, T. J., Vandjelovic, N. D., Polonenko, M. J., Li, N. R. S., Lee, A. K. C., & Maddox, R. K. (2025). The auditory brainstem response to natural speech is not affected by selective attention. PLOS Biology, 23(10), e3003407. 10.1371/journal.pbio.3003407

Teoh, E. S., Ahmed, F., & Lalor, E. C. (2022). Attention differentially affects acoustic and phonetic feature encoding in a multispeaker environment. Journal of Neuroscience, 42(4), 682–691. 10.1523/JNEUROSCI.1455-20.2021

The Joint Committee on Infant Hearing. (2019). Year 2019 Position Statement: Principles and Guidelines for Early Hearing Detection and Intervention Programs. The Journal of Early Hearing Detection and Intervention, 4(2), 1–44. doi10.15142/fptk-b748

Tremblay, K. L., Pinto, A., Fischer, M. E., Klein, B. E. K., Klein, R., Levy, S., Tweed, T. S., & Cruickshanks, K. J. (2015). Self-Reported Hearing Difficulties Among Adults With Normal Audiograms: The Beaver Dam Offspring Study. Ear and Hearing, 36(6), e290–e299. 10.1097/AUD.0000000000000195

Verhulst, S., Jagadeesh, A., Mauermann, M., & Ernst, F. (2016). Individual Differences in Auditory Brainstem Response Wave Characteristics. Trends in Hearing, 20, 2331216516672186. 10.1177/2331216516672186

